# Segregated neural explants exhibit co-oriented, asymmetric, neurite outgrowth

**DOI:** 10.1101/613703

**Authors:** David B. Pettigrew, Curtis B. Dobson, Lori G. Isaacson, Eric Leuthardt, Heather Lilley, Georgette Suidan, Keith A. Crutcher

## Abstract

Explants of embryonic chick sympathetic and sensory ganglia were found to exhibit asymmetric radial outgrowth of neurites under standard culture conditions with or without exogenous Nerve Growth Factor [NGF]. Opposing sides of an explant exhibited: a] differences in neurite length and, b] differences in terminal morphology. Strikingly, this asymmetry exhibited co-orientation among segregated, neighboring explants. The underlying mechanism[s] of the asymmetry and its co-orientation have not been discovered but appears to depend on cell clustering because dissociated sympathetic neurons do not exhibit co-orientation whereas re-aggregated clusters of cells do. This emergent behavior may be similar to the community effect described in other cell types. If a similar phenomenon exists in the embryo, or in maturity, it may contribute to the establishment of proper orientation of neurite outgrowth during development and/or injury-induced neuronal plasticity.

## Introduction

Neurons exhibit highly differentiated phenotypes and are among the most morphologically diverse cell types known. Most notable are the cellular extensions [neurites], which develop into dendrites and axons and can extend several feet in some organisms. In mammals, most neurons are also highly polarized, with the dendritic tree usually occupying a position opposite to that of the axon. When neurons are dissociated, placed into tissue culture, and given an appropriate substrate, they also extend neurites that, in some cases, mimic their phenotype in vivo, e.g., hippocampal axons and dendrites [1]. Neurons can also be cultured as tissue explants and, under permissive conditions, will exhibit a profusion of neurites that form a radial halo around the core of the explanted tissue [2].

This explant assay, initially using chick sensory or sympathetic ganglia, provided a means of detecting neurite growth-promoting substances such as Nerve Growth Factor [NGF] [2] and remains a useful bioassay to identify factors that stimulate or inhibit neurite growth in vitro. We used this explant assay over several years to detect and quantify neurite outgrowth in a variety of experimental situations [3–7]. The method, established by the early experiments of Levi-Montalcini and co-workers, involves dissection of embryonic chick sympathetic or sensory ganglia and placing small pieces [approximately 1 mm^2^] of the tissue in culture dishes coated with a suitably adhesive substrate [typically poly-ornithine] that permits attachment of the explants as well as extension of neurites. When grown in a suitable culture medium, the result is an extensive halo of neurites [8–10].

In early experiments we observed that the neurite halo often had longer neurites on one side despite the absence of any known tropic signals. Moreover, the halo was often asymmetric with differences in morphology such that one side exhibited a dense halo, often with flattened terminal endings and, at the other end, a sparser halo with thinner terminal endings. Both neurite length and morphologic asymmetry occurred under standard culture conditions with or without exogenous NGF. Examples of asymmetric neurite outgrowth have been shown in the literature but without comment [see discussion].

The asymmetric neurite outgrowth allowed for an even more striking observation. The halos from adjacent, but separate, explants exhibited co-orientation of their halos, with the longest neurites extending in the same direction. Such co-orientation was not exhibited by dissociated neurons under the same culture conditions, but was exhibited when dissociated neurons were re-aggregated, suggesting that this is an emergent property of cell aggregates.

## Materials and methods

Sympathetic chain ganglia or dorsal root ganglia [DRG] were dissected from embryonic chickens [Spafas, Boston, MA] ranging in age from E9 to E11. Following removal from the embryo, the ganglia were cut into small explants [approximately 1 mm^3^ in size] and placed in 35 mm diameter dishes [Falcon 1008, Fisher Scientific, Houston, TX] that had previously been coated overnight with poly-l-ornithine in borate buffer [pH 8.35]. Various culture media have been used but most cultures were established either in Hams F12 with or without added NGF or in serum-free Neurobasal Medium with B27 supplement [Life Technologies, Gaithersburg, MD] [11,12]. Culture durations ranged from 5 hours to 5 days. Following culture, in order to image the neurite halo, the explants were either fixed and stained for 15 minutes with a 20% silver nitrate solution for 15 minutes followed by a 10% silver nitrate solution for 5-30 minutes [as shown in figures 5A, 5B, 6-8, 10], or the living cultures were loaded with a vital dye, 15 ng/ml 5-carboxy-fluorescein diacetate AM [Molecular Probes, Eugene, OR], for 45-90 min at 37°C in Ham’s F12 medium [Sigma, St. Louis, MO] [figures 1-3, 5C, 5D, 99].

**Fig 1.**
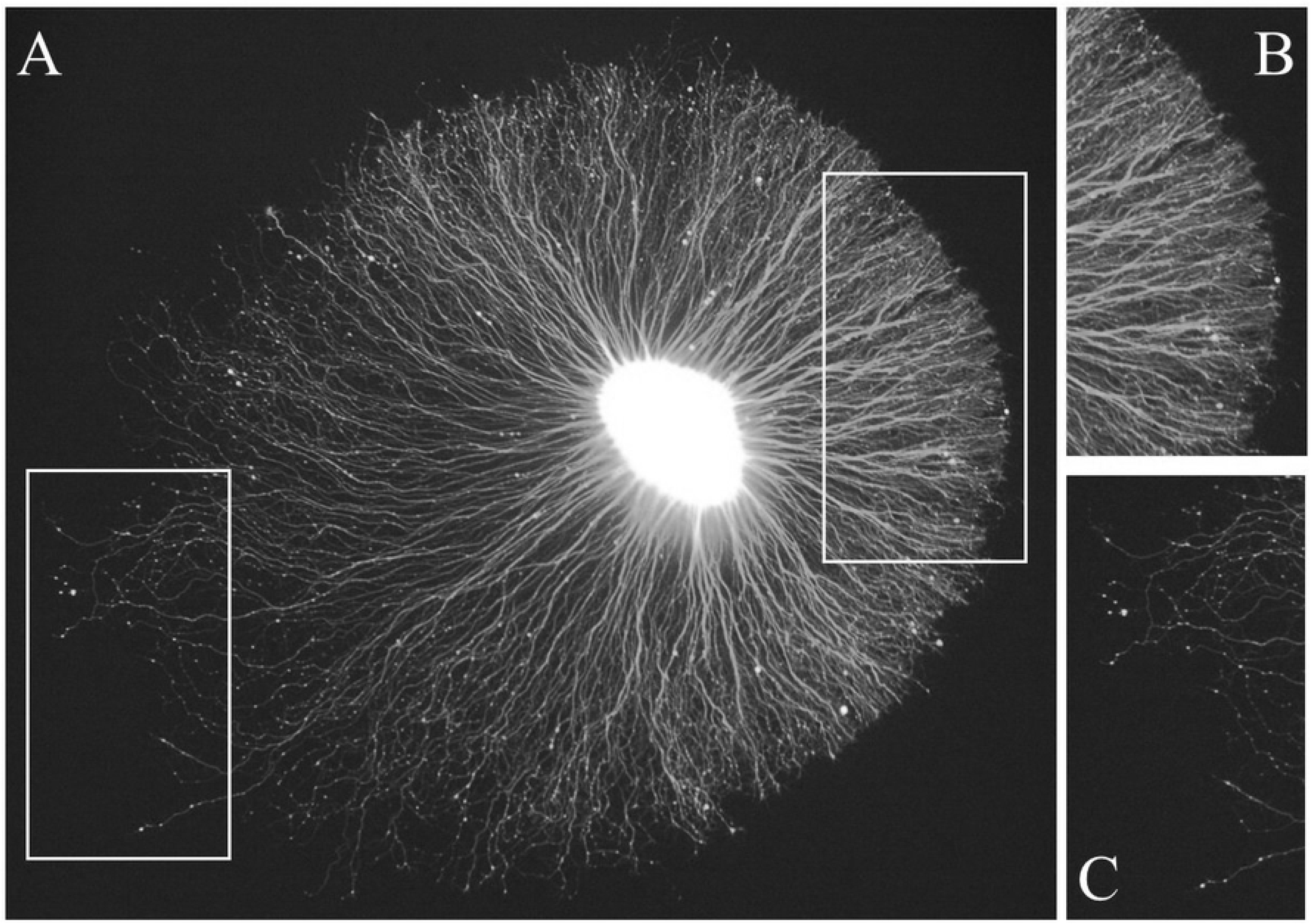
A representative example of neurite outgrowth from an embryonic chick sympathetic ganglion explant cultured in the presence of Nerve Growth Factor [NGF]. The tissue has been labeled with a vital dye to reveal the core [bright central area] and surrounding neurites [A]. The neurite halo shows asymmetric outgrowth with the shorter neurites extending to the right side where they form a well-defined edge [inset shown in panel B]. In contrast, the neurites on the left side are thinner and show more variable lengths [inset shown in panel C].

For other experiments, sympathetic neurons were dissociated by enzymatic digestion to assess outgrowth of individual neurons. Sympathetic chain ganglia were incubated with 0.25% trypsin [Sigma] for 20 min at 37°C. Trypsinization was subsequently blocked by exposure to 100% heat-inactivated fetal bovine serum [Harlan Bioproducts, Indianapolis, IN] for five minutes, and washed three times with serum-free Ham’s F12 medium [Sigma, St. Louis, MO]. The tissue was then dissociated by gentle trituration using flamed Pasteur pipettes [Fisher Scientific, Houston, TX]. In one experiment, the dissociated cells were then allowed to reaggregate before placing them in culture to assess whether dissociation would interfere with the expression of explant asymmetry and co-oriented outgrowth.

A typical explant gave rise to neurites that extended radially to form a “halo.” The resulting halo could be circular or elliptical, i.e., with a short minor axis and a longer major axis, and with the explant core occupying a position in the center of the halo or offset along the major axis [but not along the minor axis], giving rise to an asymmetric halo albeit with bilateral symmetry onon either side of the major axis. In other words, neurites systematically varied in length at different positions around the explant. In order to quantify the extent of neurite length asymmetry, a macro was developed for NIH ImageJ [source code can be provided upon request]. The perimeters of the halo and explant core were outlined and recorded by the macro. The macro fit an ellipse to the halo and identified its major axis. The macro determined the magnitude of neurite length asymmetry by measuring the distance along the major axis from the midpoint of the major axis [i.e., the center of the halo] to the explant core center. The magnitude of the neurite length asymmetry was calculated as the percent distance of the core center from the halo center to the closer end of the major axis. Thus, it was theoretically possible to have a maximum asymmetry of 100% if the entire halo extended from only one side of the explant, a phenomenon that was never observed in our cultures. In contrast, a symmetric halo, such that the explant core is directly at the midpoint of the major axis, would generate an asymmetry value of 0%.

In some cultures, there was asymmetry in growth cone morphology with little asymmetry in neurite length. In order to compare the orientation of such explants, images of individual explants were created using round image fields to eliminate cues caused by square edges. The images were then rotated randomly [each explant was rotated differently]. Two observers, blinded with respect to the original orientation, were asked to draw an equatorial line that best demarcated the morphologically distinct sides of the halo. The observers were also asked to indicate which side of the equator corresponded to the less densely organized side of the explant halo [see description in results]. The data were decoded by reversing the random rotation. The hypothesis that the explants were co-oriented was tested using the Rayleigh test [13]. Vectors corresponding to the directions of outgrowth were compared and p-values were computed based on the von Mises distribution [circular normal distribution].

## Results

The spherical core of the explant, which consists of neuronal cell bodies, the initial segments of their neurites, and Schwann cells, sometimes occupied a central location within a roughly circular and symmetric halo of outgrowing neurites. In other cases, the halo was circular but asymmetric with growth cones exhibiting morphologic differences on opposite sides of the halo. In still other cases, the halo was elliptical and asymmetric with longer neurites on one side and/or the neurites on either side of the halo exhibiting different morphologies. In cases with elliptical halos, the asymmetry was due to the explant core being off center along the major axis of the ellipse. In cases where both neurite length and morphology were asymmetric, the endings of the shorter neurites always exhibited flattened morphology and a sharply defined border.

A typical example of an explant exhibiting both types of asymmetry is shown in figure 1. The halo is wider from lower left to upper right [major axis] than from upper left to lower right [minor axis]. Along the major axis, the explant core is closer to the right side of the halo. Moreover, on the right side, the neuritic halo exhibits a highly organized distinct boundary [see discussion]. In contrast, on the left side of the halo, the boundary is less distinct and there is a gradual transition between the two sides around the halo perimeter. On the right side of the halo, the neurites are densely clustered and tightly packed [Fig 1B] whereas on the left side, the neurites tend to be longer and more variable in length [Fig 1C]. The neurites along this left boundary also appear to be thinner, and less densely organized. In some cases, single neurites extend beyond the majority of other neurites, which is rarely seen on the tightly packed side of the halo.

Another difference between the neurites extending from the right and left sides of this explant is that those on the right have extensive flattened morphologies at their tips [Fig 1B], presumably the growth cones, whereas those on the left rarely exhibit such morphologies. Many of the neurites on the left side show meandering patterns of outgrowth with narrow terminal endings [Fig 1C].

Although the example shown in figure 1 is representative of both asymmetric neurite length and morphology, there was variability in the appearance of the halos across different cultures. Some of this variability can be appreciated in figure 2, which shows examples of explants with various halo patterns. Not all explants exhibit asymmetry, either in neurite length or morphology. The example in figure 2A exhibits a circular, generally disorganized, low-density halo of neurites without obvious asymmetry or the contrasting morphologies exhibited by the explant shown in figure 1. However, even some halos with a relatively low density of neurites exhibit morphologic asymmetry, with the neurites on one side appearing thicker and with more flattened endings [Fig 2B]. Furthermore, some explants with a high density of neurites exhibit length asymmetry with little, if any, morphologic asymmetry [Figs 2C, D]. Some explants also have a few non-neuronal [presumably Schwann] cells within the immediate vicinity of the core [Fig 2D].

**Fig 2.**
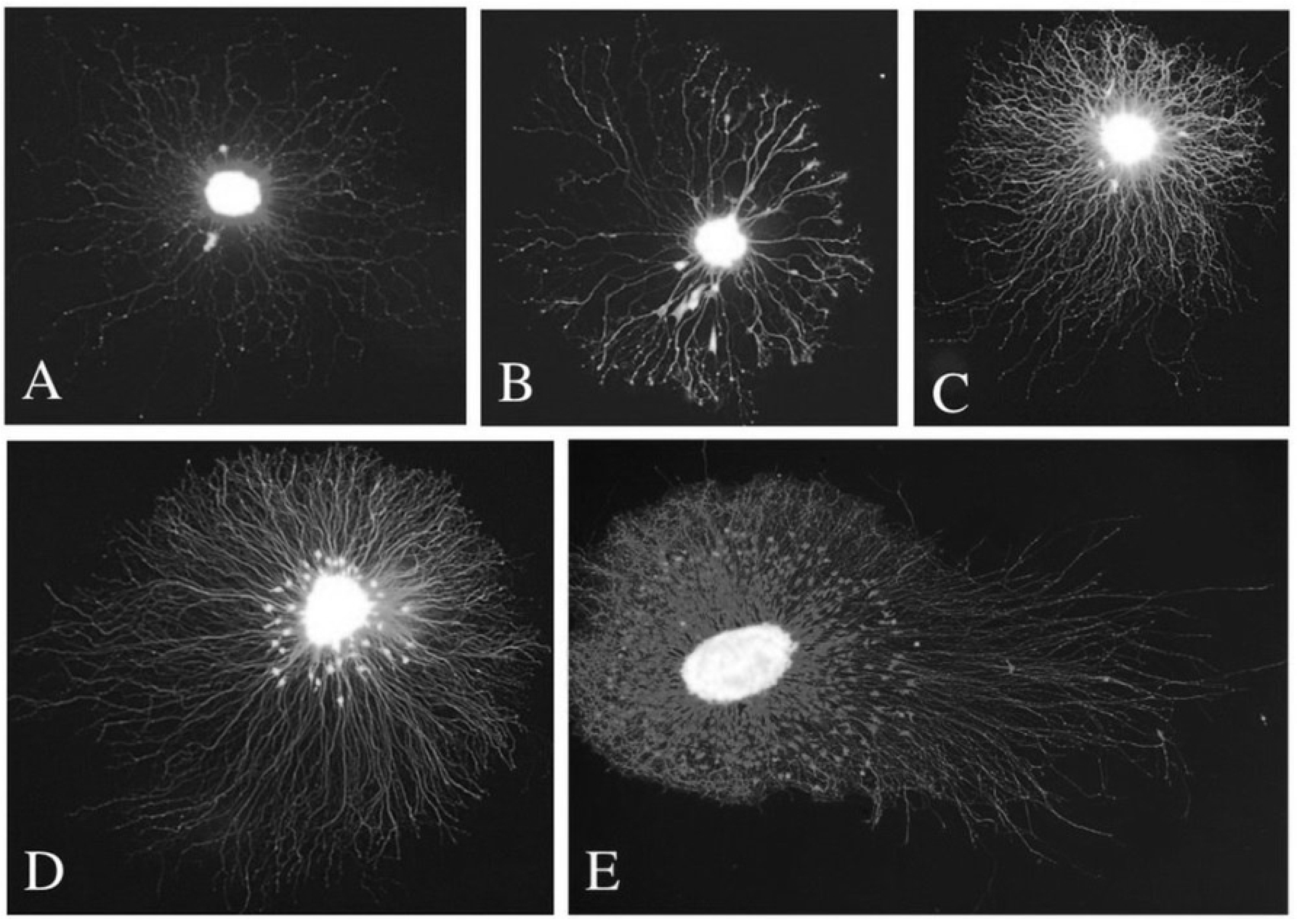
Examples of other explants with neurite halos that exhibit a range of patterns of neurite outgrowth. In some cases, there is little evidence of asymmetric outgrowth [A]. The example shown in B is an explant that has a low density of neurites but shows asymmetry in that the neurites are thicker and shorter on one side [lower right] than the other. Explants with dense halos almost always show asymmetric outgrowth, with neurites being distinctly shorter on one side compared with the other [C&D]. An embryonic chick DRG explant shows clear asymmetry with very long neurites extending from one side [E]. The other feature that distinguishes the asymmetric outgrowth of DRG explants from sympathetic ganglia is the greater extent of non-neuronal cell migration from the core of DRG explants.

The vast majority of our results are from cultures of embryonic chick sympathetic ganglia. However, we have also observed asymmetric growth in DRG explants from chick embryos of comparable age [Fig 2E]. In this example, there is a clear distinction between the opposite sides of the explant such that the neurites on the right side are much longer. The neurites on the left side are shorter and establish a sharp boundary, reminiscent of that seen in many sympathetic explants. The main difference between the pattern of outgrowth from sympathetic and DRG explants is that the DRG explants have an extensive halo of non-neuronal cells that almost reaches to the limit of the shorter neurite border [compare Figs 2D and 2E]. However, unlike the neurite outgrowth, the distribution of these non-neuronal cells is not notably asymmetric.

Because most of our cultures were established with sympathetic explants, we analyzed the prevalence and extent of outgrowth asymmetry with this tissue. Neurite length asymmetry was measured using a customized NIH Image J macro to define the relationship between the explant core and its corresponding halo as described above [Fig 3A-C]. The halo perimeter exhibits an irregular border, especially in the region where neurite length is most variable. Outgrowth asymmetry was expressed as the percent distance of the center of the explant core from the center of the major axis of the halo. In the example shown [Fig 3A-C] asymmetry was estimated at 33.8% [i.e., the explant core is displaced 33.8% towards the lower right side of the halo]. Examples of the extent of length asymmetry calculated in this manner are shown in figure 3D-F. The extent of asymmetry measured in this way showed examples with virtually no length asymmetry [Fig 3D, 1.8%], where the explant halo is virtually symmetric, and other cases in which there is greater asymmetry [Fig 3E, 19.2%] and [Fig 3F, 28.7%].

**Fig 3.**
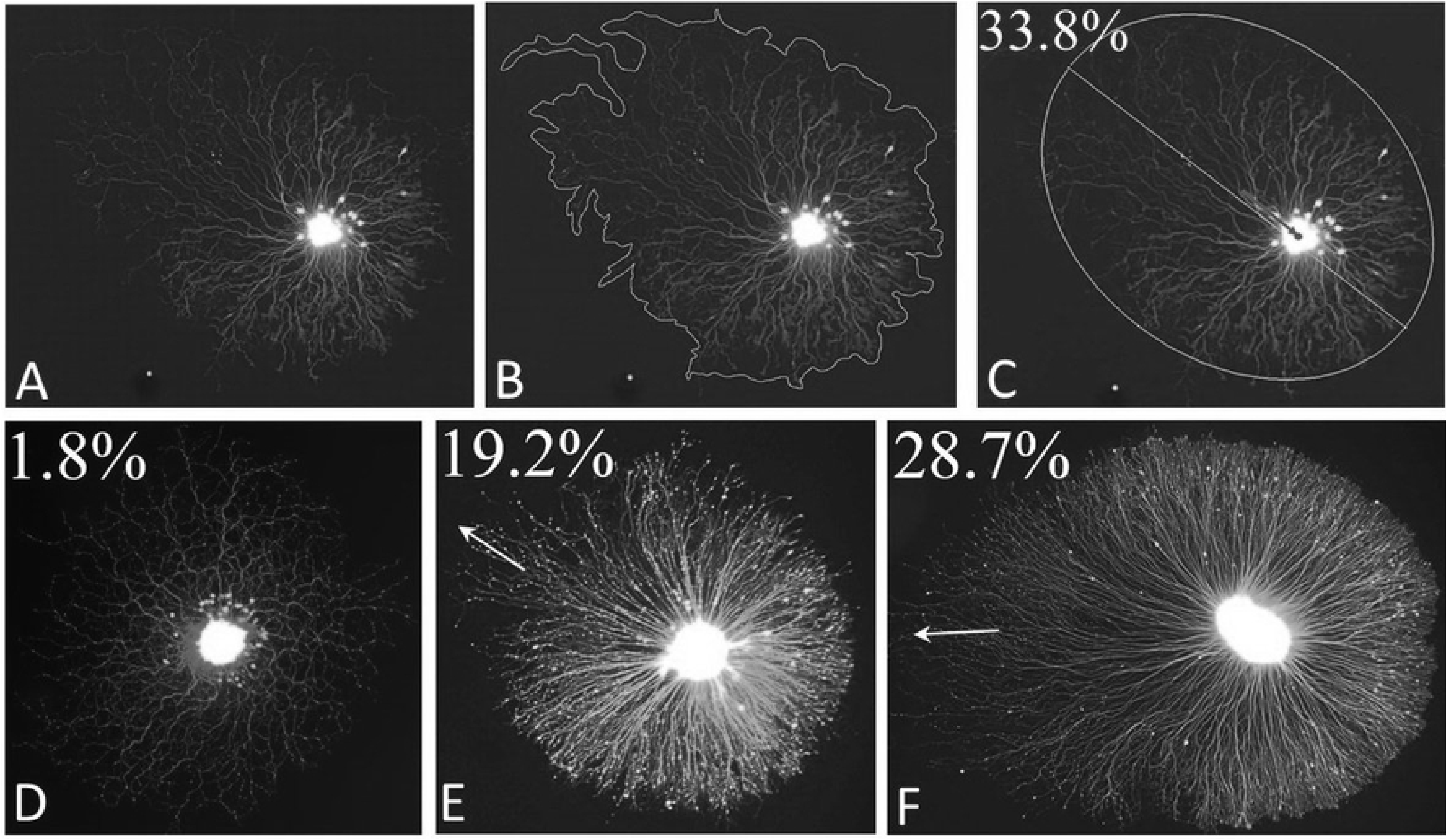
Quantification of the degree of neurite length asymmetry and its orientation. An example of the method is shown using the explant in panel A. A contour was drawn around the explant halo [B] and explant core [not shown]. Using the “Fit Ellipse” tool, ellipses were fit to the halo [C] and the core [not shown]. The centers and major axis lengths of both ellipses were identified. The center of the explant core was compared to the distance from halo center to nearest end of the major axis [arrow in C]. The magnitude of neurite length asymmetry was taken to be the percent displacement of the core center from the halo center to the nearest end of the major axis [in this case, 33.8%]. Examples of other explants analyzed in this way are shown in D-F.

To obtain some indication of the prevalence and extent of neurite length asymmetry, 603 explants from thirteen cultures were analyzed according to the procedure described above. As shown in the histogram in figure 4A, explants showed asymmetry ranging from 0 to 60% with the mode of the distribution at around 35%. Of these 603 explants, 440 of the explants [73%] exhibited an asymmetry of at least 15%. The extent of asymmetry did not correlate with either core area [Fig 4B] or halo area [Fig 4C].

**Fig 4.**
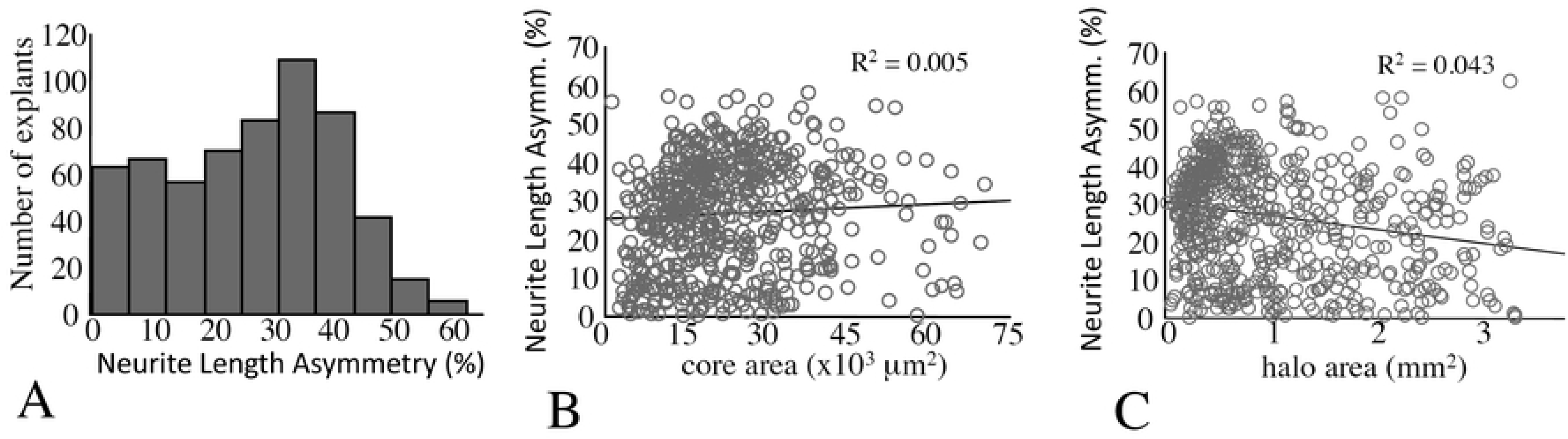
Prevalence and extent of neurite length asymmetry. [A] Histogram showing the prevalence of neurite length asymmetry in 603 explants. Bivariate scattergrams showing the distribution of asymmetry as a function of explant core [B] and halo [C] areas. The extent of neurite length asymmetry showed no direct correlation with either variable.

**Fig 5.**
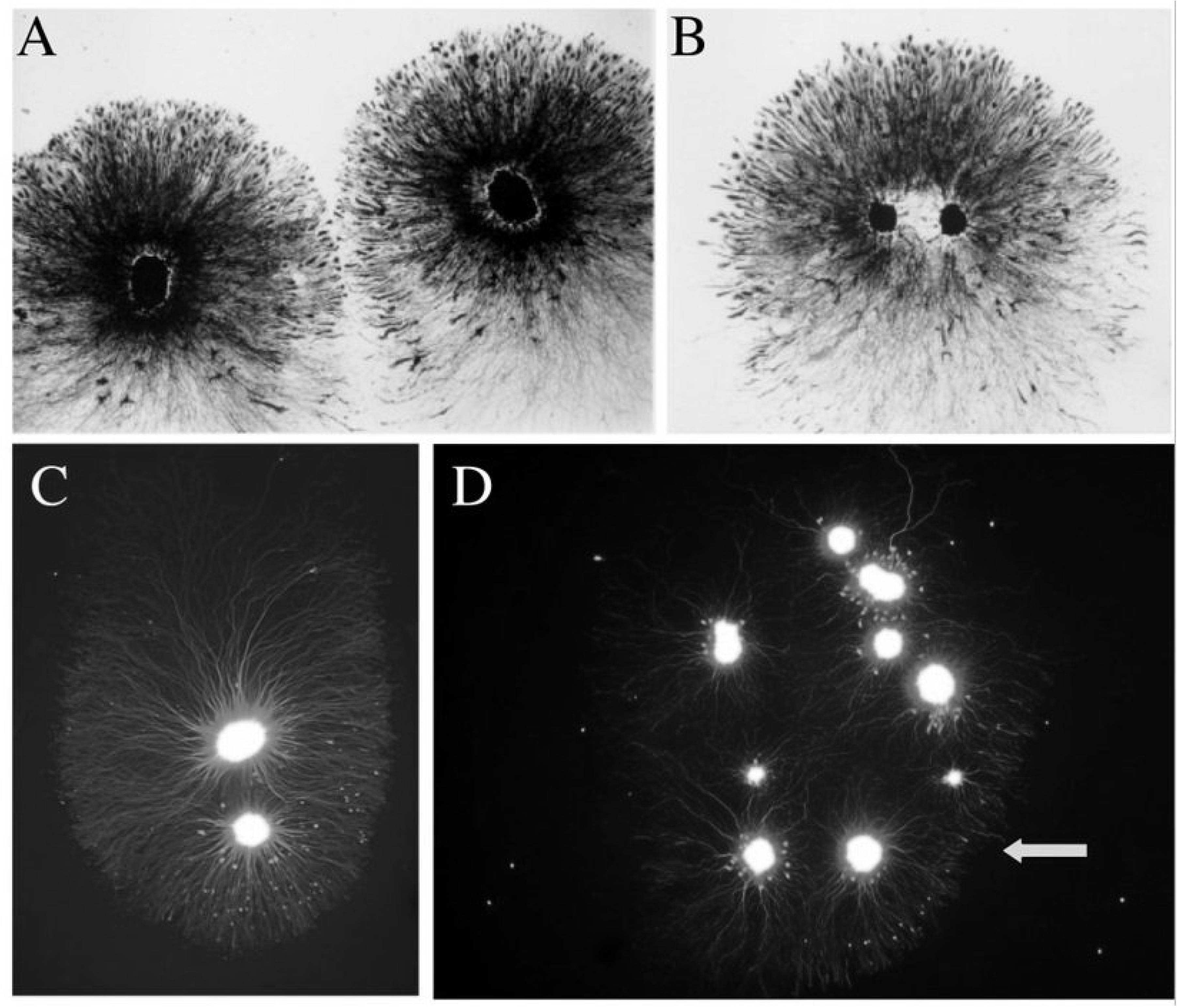
Explants with shared halos. Two explants growing close to each other have extended asymmetric halos, each of which shows similar orientation [A]. Two explants may contribute to a common halo that shows asymmetry with the cores situated side-by-side [B] or in “tandem” [C]. A collection of nine explants has extended halos that collectively exhibit a front of organized neurites [arrow] on one side that are shorter than those on the opposite side of the collective halo [D].

In addition to the common occurrence of outgrowth asymmetry, a second higher order level of coordination was observed when multiple explants were cultured in the same dish. In these cases, two adjacent explants usually showed similar orientation of asymmetry [Fig 5A]. If two explant cores were in sufficient proximity to each other, they would share a common halo that also exhibited asymmetry [Figs 5B, C]. In rare cases, more than two cores contributed to a common halo that was asymmetric [Fig 5D]. In cases where explants were segregated, their asymmetric halos exhibited co-orientation with neighboring explants. This was best observed with silver staining because of the ability to visualize several explants with low power magnification using bright field illumination. An example of a field of eight segregated sympathetic explants in a single dish is shown in figure 6. Each shows evidence of asymmetric neurite outgrowth, in neurite length, density and terminal morphology. In addition, the orientation of the longest neurites in each halo is in the same direction, in this case toward the right side of the field. None of the explants are in physical contact with each other. Furthermore, co-orientation was a local phenomenon, i.e., explants on the other side of the dish or in adjacent dishes did not necessarily have the same orientation as each other.

**Fig 6.**
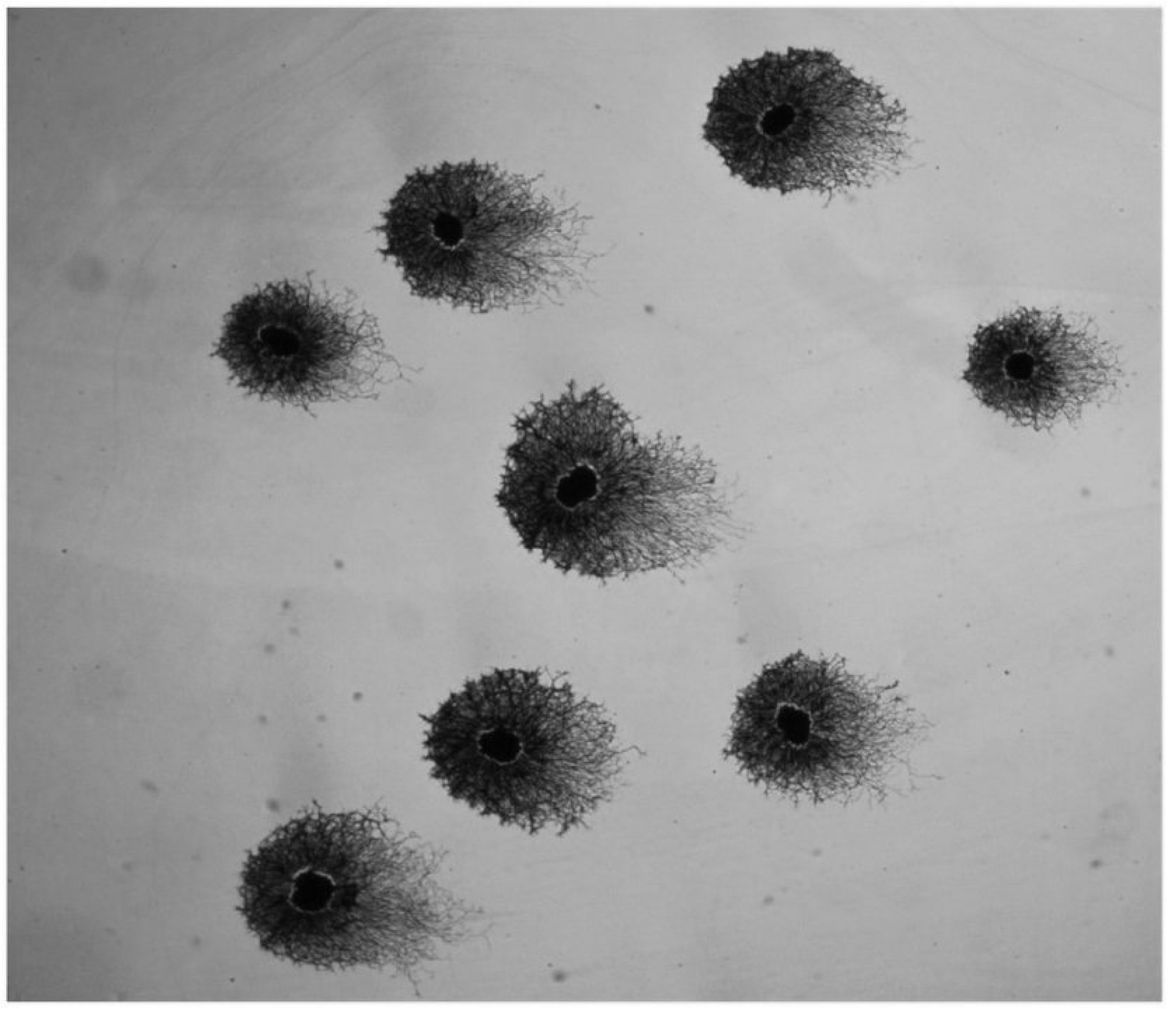
A field of silver-stained sympathetic explants exhibiting co orientation of neurite halos. Each of the eight explants shown here exhibits an asymmetric halo and the longest neurites within each halo extend to the right side of the field in each case. Such co-orientation of halos was a common occurrence among neighboring explants although there was no consistent orientation from one culture dish to another.

Another example of a culture in which several explants show co-orientation is shown in figure 7. In this example, there was little asymmetry in neurite length [although the silver stain does not reveal the longest individual neurites at this magnification] although there does appear to be asymmetry in growth cone morphology. In order to evaluate the extent of co-orientation, the six explants in this field were evaluated by two subjects blinded to the original orientation who calculated a vector for each explant using randomly rotated images [see methods]. Rayleigh analysis of the polar plot of these averaged vectors demonstrated statistically significant co-orientation [r = 0.998; *p* < 0.004; Fig, 7, inset].

**Fig 7.**
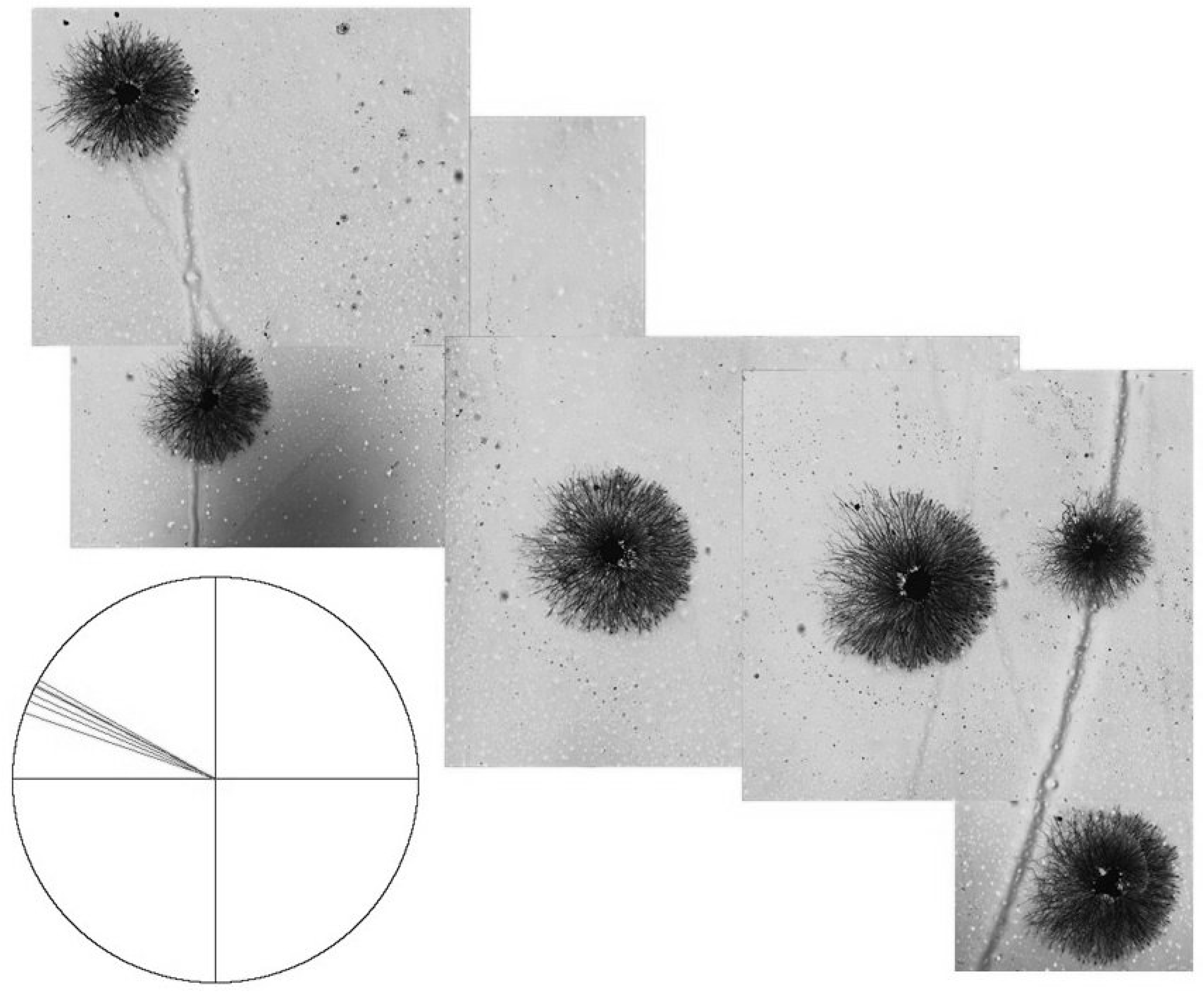
Six silver stained explants with cores centered within circular neurite halos. Morphologic differences in the halo are evident, with tightly packed, well organized fronts of neurites to the right and less densely packed, less organized fronts of neurites to the left. Vector diagram [inset] shows these halos are highly co-oriented based on blinded assessments of halo asymmetry [Rayleigh r = 0.998; p < 0.004].

A few cultures of DRG explants were also established to determine whether co-oriented asymmetry also occurs with this neural tissue. Figure 8 shows an example of a culture in which four DRG explants exhibit such co-orientation. The longest neurites are all generally oriented in the same direction for each explant. Not shown is another explant that was situated approximately four millimeters away that showed the same orientation as those in the cluster.

**Fig 8.**
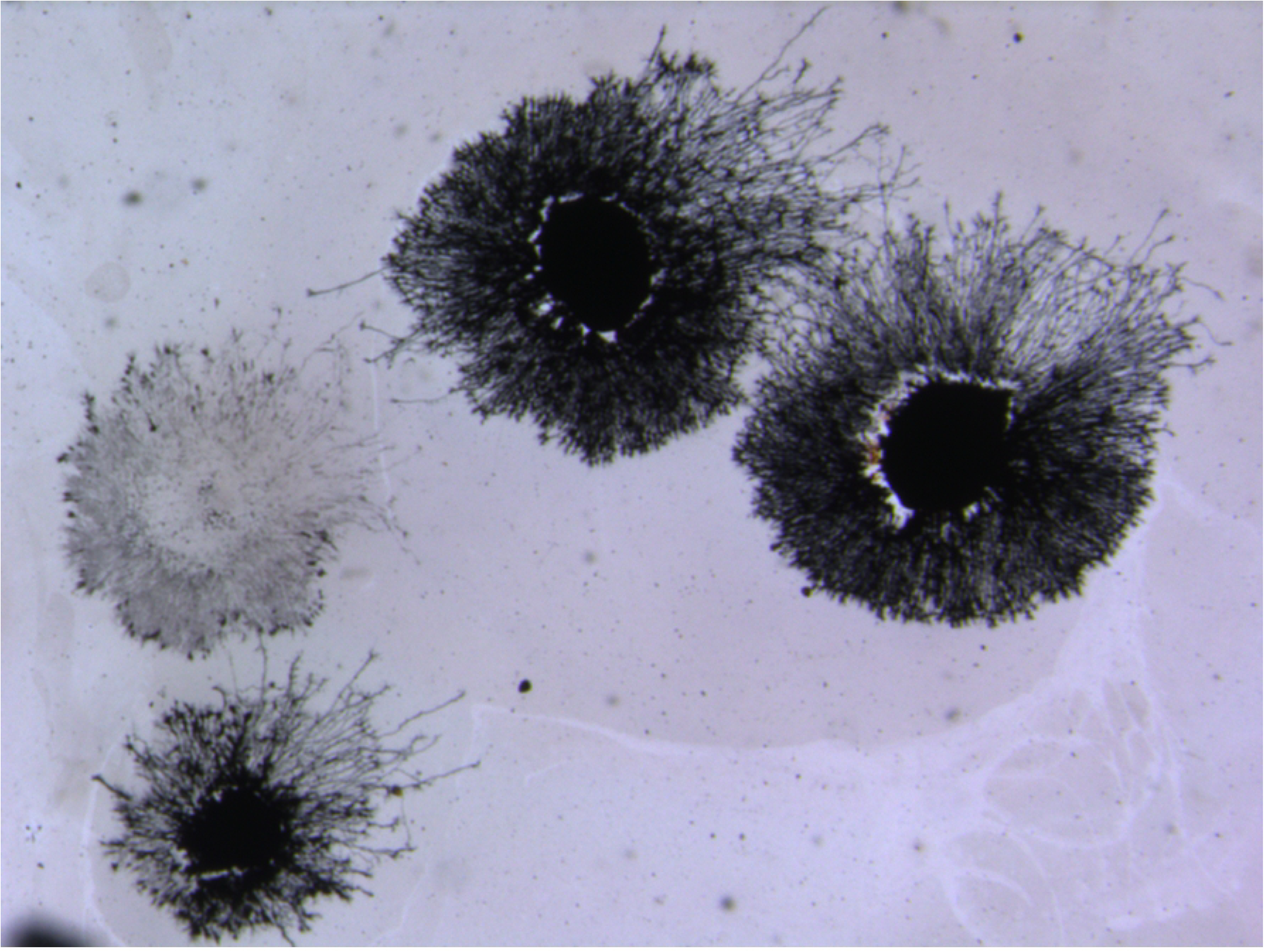
A cluster of four dorsal root ganglia explants exhibiting co-oriented asymmetric neurite outgrowth. The longest neurites extend to the upper right. The upper left explant was lost during the staining procedure but enough of the halo is present to appreciate the asymmetric orientation of the neurites.

To determine whether the orientation of explant halos was also reflected in the orientation of neurites from individual cells in the same cultures, we combined dissociated cells with explant cultures [figure 9]. In spite of the clear asymmetry in the neurite halo of the explant, there was no apparent directional orientation of outgrowth from individual neurons. Finally, to assess the possibility that the asymmetry and co-orientation depend on cues retained ex vivo after being established in the embryo, we dissociated ganglia and then allowed the cells to re-aggregate before placing them in culture. The re-aggregated explants also showed asymmetric co-oriented outgrowth such that the longest neurites extended in the same direction [arrows in figure 10].

**Fig 9.**
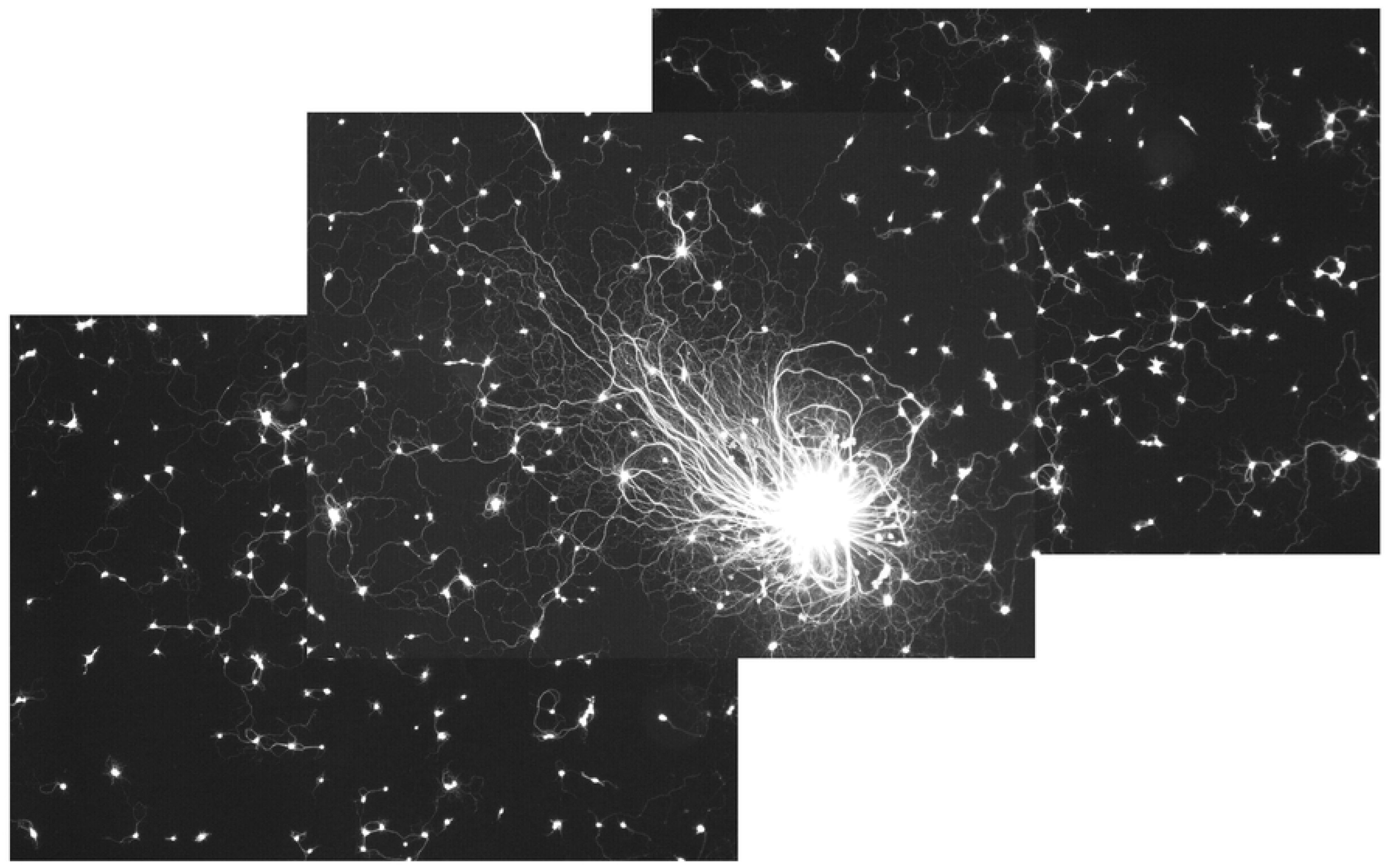
A sympathetic explant with asymmetric outgrowth. The longest neurites extend towards the upper left of the field. This explant was co-cultured with dissociated neurons, which show no consistent orientation in outgrowth.

**Fig 10.**
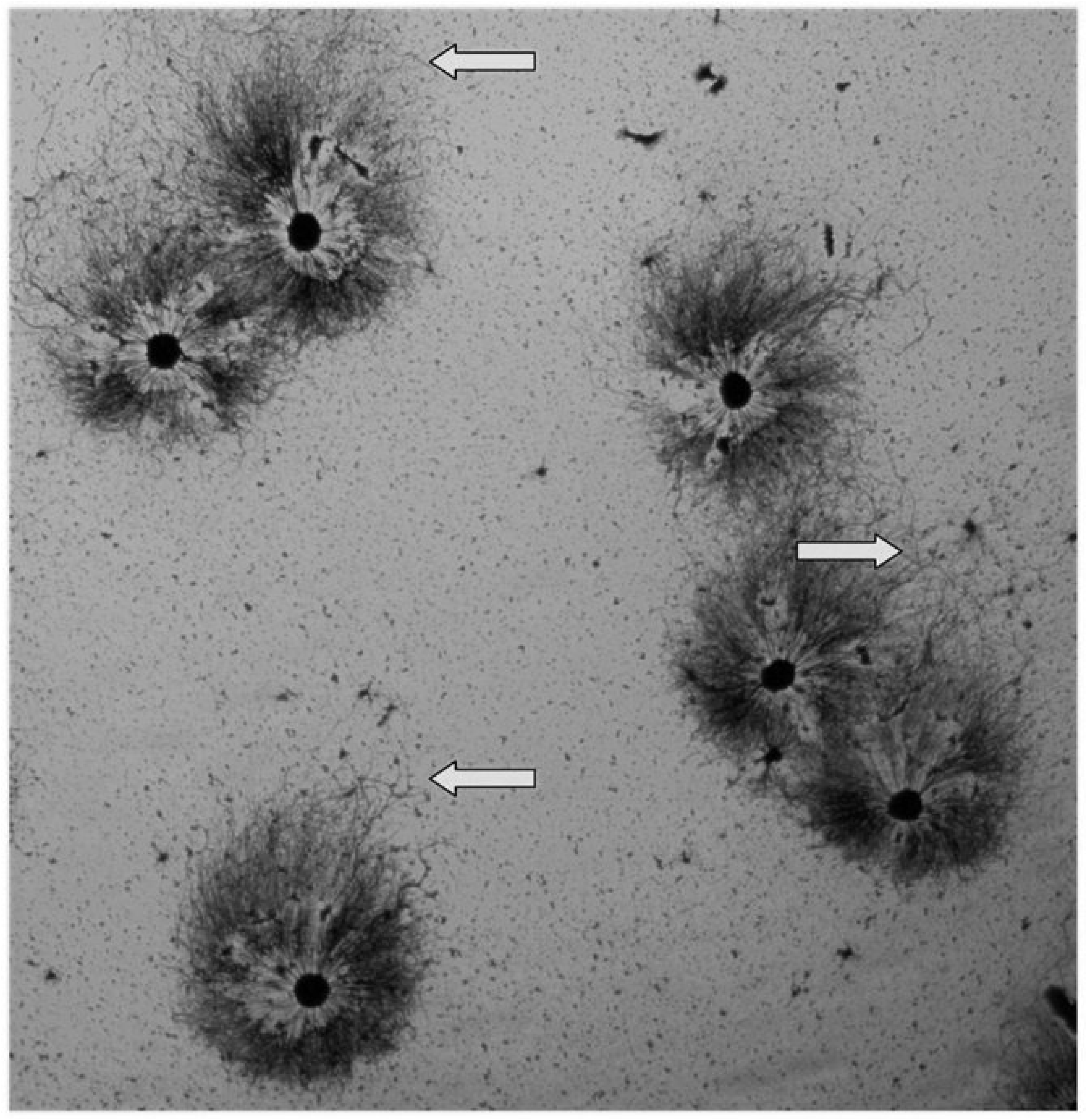
A field of “reaggregated” explants is shown stained with silver nitrate. The ganglia were first dissociated and the cells were then allowed to form clusters before plating. The resulting neurite halos exhibit asymmetry and the longest neurites extend in the same general direction [towards the top of the field, as indicated by the arrows]. The neurites on the opposite side tend to be shorter and to establish a sharper boundary.

## Discussion

This study demonstrates a seemingly simple phenomenon whereby neural tissue [explants of embryonic chick sympathetic or sensory ganglia] undergo asymmetric neurite outgrowth in nominally uniform growth environments. Furthermore, this asymmetric growth shows regional similarity and is observed with cell aggregates but not with single cells. This emergent asymmetry supports the notion that there is a tissue-level mechanism that serves to organize neural tissue. This behavior could have fundamental implications for the mechanisms underlying the transition from single cells to more complex multicellular structures in both ontogeny and phylogeny.

The neurite halo assay was first developed to detect neurite growth-promoting substances released by tumor tissue [2,14,15] and has been used by numerous investigators. The original description of the results obtained when neural tissue was co-cultured with tumor tissue bears similarity to our results. The neurites from a DRG explant facing the tumor were described as showing “*maximal density and a very straight course*” whereas those on the opposite side were described as “*less dense and longer; the direction of their outgrowth is less straight, and some take more winding routes”* [2]. The majority of explants shown in most of the early publications using this method do not include the full explant halo so it is difficult to appreciate the asymmetry being described. However, a number of images from this original work can be found, including one on the cover of a book published by Levi-Montalcini clearly showing asymmetry [16]. The same example was used in a review paper [17]. In fact, the description of halo asymmetry in the 1954 paper cited above could well apply to the example shown in our figure 1. The main experimental difference being that our neural explants were never cultured in the presence of non-neural tissue. Nor did we observe explant halos oriented towards each other.

The shape of the neuritic halo from “control” explants was described by Levi-Montalcini as being a “*circular or ellipsoidal, perfectly geometrical ring around the explant”*[18]. Although an ellipse can theoretically be defined with the explant core at its center, the examples illustrated by Levi-Montalcini and colleagues [for example, figure 3B in [15] as well as our results], almost always have the core nearer to one focus of an ellipse. Furthermore, the long trail of neurites often present on one side gives the explant the appearance of a tear-drop or comet-shaped halo. Of the published images in the literature, some show mainly radially-symmetric outgrowth [9,10] but there are numerous examples of asymmetric halos as well, e.g., figure 4b of [19], figure 1b in [20], figure 2 in [21], figure 7 in [22], figure 2A in [23], and figure 1 in [24]. However, many published images show only a small portion of the explant and/or its halo so that it is not possible to determine whether asymmetric outgrowth occurred.

As in the original work involving co-cultures of neural tissue with non-neural tissue, preferential outgrowth from neural explants co-cultured with other tissue types has been documented [25–31]. Other studies have used orienting stimuli, such as a local source of a neurotrophic factor [9] or application of an electric field [32,33] to elicit asymmetric outgrowth. In most such studies, there is little information provided on the prevalence of asymmetry in control cultures.

Unlike other studies where asymmetric outgrowth was reported in the presence of other tissue or some other stimulus, the explants in the present study were cultured in nominally homogeneous conditions. Although not all explants showed asymmetry, neurite outgrowth from embryonic chick sympathetic ganglia was at least 15% asymmetric in 73% of the explants quantified [the mode is 35% elliptical asymmetry, approximately that shown in the example shown in figure 3C]. Asymmetric outgrowth did not depend on the presence of exogenous NGF as long as other conditions permitted the establishment of a neurite halo, e.g., the use of NeuroBasal medium. Asymmetry in neurite length and morphology did not always occur together [Fig 7] but when they did, the shorter neurites were inevitably those with more flattened terminals, presumably growth cones.

The mechanism[s] underlying the asymmetry or co-orientation is uncertain. Detection of co-orientation depends on the presence of halo asymmetry. In fact, it was the co-orientation that first brought our attention to the asymmetric outgrowth. However, whether a common mechanism accounts for both phenomena is not clear. Although individual neurons exhibited extensive neurite outgrowth under the same culture conditions, co-orientated growth was only observed in explants or re-aggregated cell clusters. This suggests that co-oriented outgrowth is an emergent property requiring a minimal number of cells analogous to what has been termed the “community effect” in other systems, such as the differentiation of muscle cells [34–40].

In light of the evidence that various stimuli can influence neurite outgrowth in culture, including diffusible growth factors, substrate-bound factors, and bioelectric phenomena, a number of hypotheses can be envisioned. We have no direct evidence to distinguish these, although the fact that the orientation of outgrowth is not consistent from dish to dish in the same experiment argues against a global field effect such as gravity.

If the in vitro observations reported here are at least partly attributable to phenomena that occur during development, further studies of explants cultured under similar conditions could reveal relevant mechanisms also operating in vivo. At the very least, the co-oriented asymmetric outgrowth reported here suggests that there are tissue-level mechanisms that serve to organize neural tissue in a way not previously reported. Additional experiments will be required to account for the mechanism underlying both the expression of asymmetry and the co-orientation of neuritic halos.

## Acknowledgements

Numerous individuals participated in experiments in which the phenomenon of asymmetric outgrowth was observed. The help and participation of Jill Sigmon, Jean Weingartner, Kristina Bielewicz, Lori Gulley, Sarah Anthony, Billy Toro, and Alex McCullough are gratefully acknowledged. The insightful comments and suggestions of Dr. Stephen Robinson, Dr. Ron Huston, Dr. Monica Brauer, and Dr. John Quinlan have also been extremely helpful.

